# Understanding hepatitis B virus dynamics and the antiviral effect of interferon-α treatment in humanized chimeric mice

**DOI:** 10.1101/2020.07.28.224840

**Authors:** Laetitia Canini, Yuji Ishida, Masataka Tsuge, Karina Durso-Cain, Tje Lin Chung, Chise Tateno, Alan S. Perelson, Susan L. Uprichard, Kazuaki Chayama, Harel Dahari

## Abstract

**Background:** Whereas the mode of action of lamivudine (LAM) against hepatitis B virus (HBV) is well established, the inhibition mechanism(s) of interferon-α are less completely defined. To advance our understanding, we mathematically modelled HBV kinetics during pegylated interferon-α-2a (pegIFN), LAM and pegIFN+LAM treatment of chronically HBV-infected humanized uPA/SCID chimeric mice.

**Methods:** Thirty-nine uPA/SCID mice with humanized livers whose pre-treatment steady-state serum HBV reached 9.2±0.4 logIU/mL were treated with pegIFN, LAM or pegIFN+LAM for 14 days. Serum HBV DNA and intracellular HBV DNA were measured frequently. We developed a nonlinear mixed effect viral kinetic model and simultaneously fit it to the serum and intracellular HBV DNA data.

**Results:** Unexpectedly, even in the absence of an adaptive-immune response, a biphasic decline in serum HBV DNA and intracellular HBV DNA was observed in response to all treatments. Modeling predicts that the first phase represents pegIFN inhibiting intracellular HBV DNA synthesis with efficacy of 86%, which was similar under LAM and pegIFN+LAM. In contrast, there were distinct differences in HBV decline during the 2nd phase which was accounted for in the model by a time-dependent inhibition of intracellular HBV DNA synthesis with the steepest decline observed during pegIFN+LAM (0.46/d) and the slowest (0.052/d) during pegIFN mono-treatment.

**Conclusions:** Reminiscent of observations in patients treated with pegIFN and/or LAM, a biphasic HBV decline was observed in treated humanized mice in the absence of adaptive immune response. Interestingly, combination treatment does not increase the initial inhibition of HBV production; however, enhancement of second phase decline is observed providing insight into the dynamics of HBV treatment response and the mode of action of interferon-α against HBV.

**Author Summary:** Chronic hepatitis B virus (HBV) infection remains a global health care problem as we lack sufficient curative treatment options. Elucidating the dynamic of HBV infection and treatment at the molecular level would potentially facilitate the development of novel, more effective HBV antivirals. Currently, the only well-established small animal HBV infection model available is the chimeric uPA/SCID mice with humanized livers; however, the HBV infection kinetics under interferon-α (IFN) in this model system have not been determined in sufficient detail to support the in-depth studies of HBV treatment response needed to identify/confirm more effective drug targets. In this study 39 chronic HBV-infected uPA/SCID humanized mice treated with IFN and/or lamivudine were analysed using a mathematical modelling approach. We found that IFN main mode of action is blocking HBV DNA synthesis and that 73% of synthesized HBV DNA per are secreted from infected cells. Our data-driven mathematical modeling study provides novel insights into IFN anti-HBV mechanism(s) and viral-host interplay at the molecular level.

## Introduction

Despite the availability of an effective vaccine since 1981, hepatitis B virus (HBV) continues to impose an enormous global health burden. Over 260 million people worldwide are chronically HBV infected, causing persistent hepatitis and hepatocellular carcinoma (1, 2). While currently available antiviral drugs can suppress virus replication, less than 10% of patients treated with interferon-based therapies and less than 1% treated long-term with nucleos(t)ide analogues (NUCs) exhibit clearance of the virus and/or clearance of serum hepatitis B surface antigen (HBsAg) by HBs antibodies (termed HBsAg seroconversion), the latter being clinically defined as “functional cure” of the infection (3–6). Thus, the development of more effective/curative therapeutics is urgently needed.

One way to facilitate the development of better antiviral strategies is to obtain a deeper understanding of HBV infection dynamics. Two studies monitoring HBV DNA in the serum of treated patients reported that pegylated interferon-α2a (pegIFN) monotherapy was less effective in lowering viral load than lamivudine (LAM) monotherapy with the highest response seen under pegIFN+LAM combination treatment (6, 7). Early viral kinetic analysis and mathematical modeling of these pegIFN, LAM and pegIFN+LAM treatments indicated that serum HBV DNA typically dropped in a biphasic manner consisting of a rapid 1^st^ phase that was predicted to depend on the viral half-life and drug efficacy in blocking viral production, followed by a 2^nd^ slower phase that was projected to represent the loss rate of infected cells (8). However, since it is not feasible to obtain liver biopsies from patients during antiviral treatment the nature of the observed biphasic viral decline in patients’ serum is unknown.

In this study we investigated, for the first time, both serum and intracellular HBV viral kinetics in 39 uPA/SCID chimeric humanized mice undergoing pegIFN, LAM or pegIFN+LAM treatment, and developed a mathematical model to provide insights into HBV dynamics and pegIFN mechanism of action (MOA).

## Methods & Material

### Production of uPA/SCID chimeric mice with humanized liver

Thirty-nine uPA^+/+^/SCID^+/+^ chimeric mice with humanized liver were produced as described previously (9–11). Three different commercially available frozen lots of human hepatocytes (2 year old Hispanic male, 2YM; 5 year old African American male, 5YM; 1 year old Caucasian female, 1YF) were purchased from BD Biosciences Discovery Labware, San Jose, CA, USA (2YM and 5YM) and In Vitro Technologies, Baltimore, MD, USA (1YF). 2YM donor had IL28B, rs12979860, CC genotype, 5YM had IL28B TT genotype, and 1YF IL28B genotype was unknown. They were thawed and 2.5×10^5^ viable hepatocytes were transplanted into 2- to 4-week-old uPA/SCID mice via intrasplenic injection. All animal protocols described in this study were performed in accordance with the Guide for the Care and Use of Laboratory Animals and approved by the Animal Welfare Committee of Phoenix Bio Co., Ltd.

### Serum human albumin (hAlb) quantification

Because hAlb concentration in the serum correlates with the degree of human hepatocyte repopulation observed by immunohistochemical staining of liver sections with human specific antibodies (10), hAlb concentration in mouse serum was measured by latex agglutination immunonephelometry (LX Reagent “Eiken” Alb II; Eiken Chemical Co., Ltd., Tokyo, Japan) at 3 and 6 weeks after hepatocyte transplantation, and thereafter once a week until treatment. These hAlb levels were used to estimate the replacement index of human hepatocytes in individual mice as previously described (10). Levels of human serum albumin were then also measured at baseline and days 3, 7, 10 and 14 during treatment to monitor for changes in human hepatocyte number/function.

### Infection of uPA/SCID humanized mice with HBV

The original virus (HBV genotype C, from an HBeAg-positive patient), provided by Dr. Sugiyama (12), was used to create the viral stock needed for these experiments by amplifying the virus in uPA/SCID chimeric mice. Specifically, after observing stable high-level HBV viremia in inoculated mice, sera was collected and pooled. HBV DNA level was determined by quantitative real-time PCR as reported previously (13). For experiments, pooled serum containing 10^4^ copies of HBV DNA (i.e. 9740 IU) was intravenously injected into chimeric mice whose hAlb levels were 4.2-15.0 mg/mL by 7 to 11 weeks after human hepatocyte transplantation (corresponding to a replacement index of 53.7-89.3%). All 39 mice were reached high HBV steady state levels as described previously (11).

### Antiviral drug treatment

Lamivudine (LAM) was purchased from Tokyo Chemical Industry Co., Ltd. (Tokyo, Japan) and pegylated interferon α-2a (pegIFN) was purchased from Chugai Pharmaceutical Co., Ltd. (Tokyo, Japan). Antiviral treatment was started once all mice had achieved steady state serum HBV DNA levels at 9 weeks post-HBV inoculation and continued for 2 weeks. Three therapy protocols were followed. Lamivudine (30 mg/kg) was given orally every day alone, pegIFN (30 μg/kg) was injected subcutaneously twice weekly alone, or both drugs (pegIFN+LAM) were given. Nine chimeric mice (3 mice transplanted with donor 5YM hepatocytes and 6 mice transplanted with donor 2YF hepatocytes) were treated with pegIFN monotherapy. Twenty-one mice were treated with LAM only. Of these, 9 mice were transplanted with donor 2YM hepatocytes, 4 mice with donor 1YF, and 8 mice with donor 5YM. Nine chimeric mice (3 with 5YM and 6 with 2YF) were treated with pegIFN+LAM combination therapy (**Supplementary Table 1**).

### Intracellular HBV DNA

Extraction and quantification of intracellular HBV DNA were performed as recently described (11). The intracellular viral loads were measured for 5 mice sacrificed prior to treatment to determine pre-treatment/steady state intracellular HBV DNA levels. Nine mice were sacrificed at day 3 post-treatment (3 from each treatment group) and 30 mice were sacrificed at day 14 post-treatment (6, 18 and 6 from pegIFN, LAM and pegIFN+LAM treatment groups, respectively) (**Supplementary Table 1**). Because fitting the model to the viral kinetics in individual animals requires longitudinal intracellular data for each mouse, but intracellular analysis can only occur at one time point (i.e. the time of sacrifice), sampling from a uniform distribution of the measurements from the other mice of the same treatment group (days 3 and 14) or at baseline (day 0) was performed in order to estimate the other intracellular HBV DNA levels at the other time points for the sacrificed mice and thus supplement the intracellular longitudinal data.

### Statistical analysis

Data is expressed as arithmetic mean ± standard deviation or median value with range. Non-parametric tests included the Kruskal-Wallis test and Friedman test for repeated measurements. Multiple comparisons (pairwise by Conover-Iman transformation) were corrected by the Bonferroni-Holm method. Bivariate correlations were quantified by Spearman rank-correlation coefficients. Results with p-values ≤ 0.05 were considered statistically significant. Serum hAlb slopes were estimated by linear regression. Statistical analyses and graphs were computed with BiAS (version 10.03), R (version 2.15.1) and Microsoft Excel (version 14.2.4). Serum hAlb kinetics showed mono- or biphasic decline. In consequence, it was modelled by a single slope, s1, for monophasic decline or a segmented linear regression with two slopes, s1 and s2, and a breakpoint at t1 for biphasic decline.

### Mathematical modeling

We developed a multicompartmental model (**Equation 1** and **Figure 1**), consisting of infected human hepatocytes, *I*, free circulating virions, *V*, and total intracellular HBV DNA, *D*. Since we recently demonstrated, by immunostaining, that in uPA/SCID chimeric mice nearly all human hepatocytes (~95%) are infected by 8-10 weeks after HBV inoculation (11), no availability of uninfected human hepatocytes was assumed. Model equations are as follows

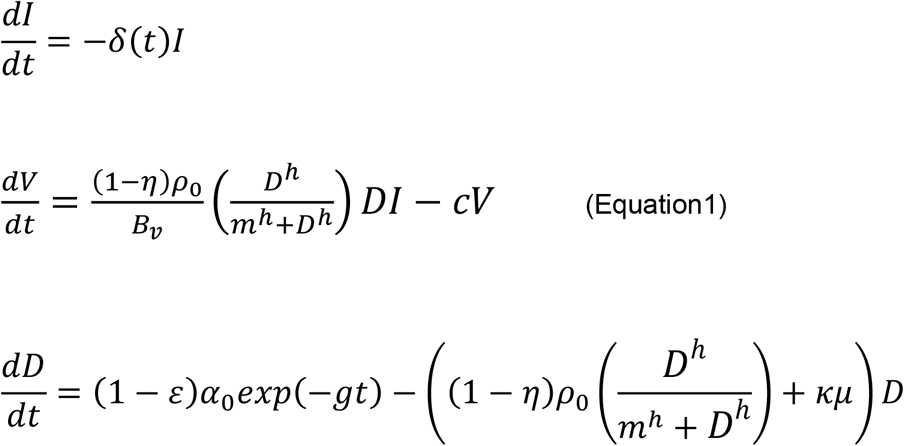

where total intracellular HBV DNA, *D*, is produced at rate *α*_0_, and lost due to degradation with rate constant *μ*. For simplicity, we do not distinguish between multiple intracellular replication steps such as the generation of viral RNA from covalently closed circular (cccDNA) within the nucleus, cytoplasmic capsid assembly, or generation of HBV single-stranded DNA by reverse transcription and maturation to double-stranded DNA inside the capsid.

**Figure 1:**
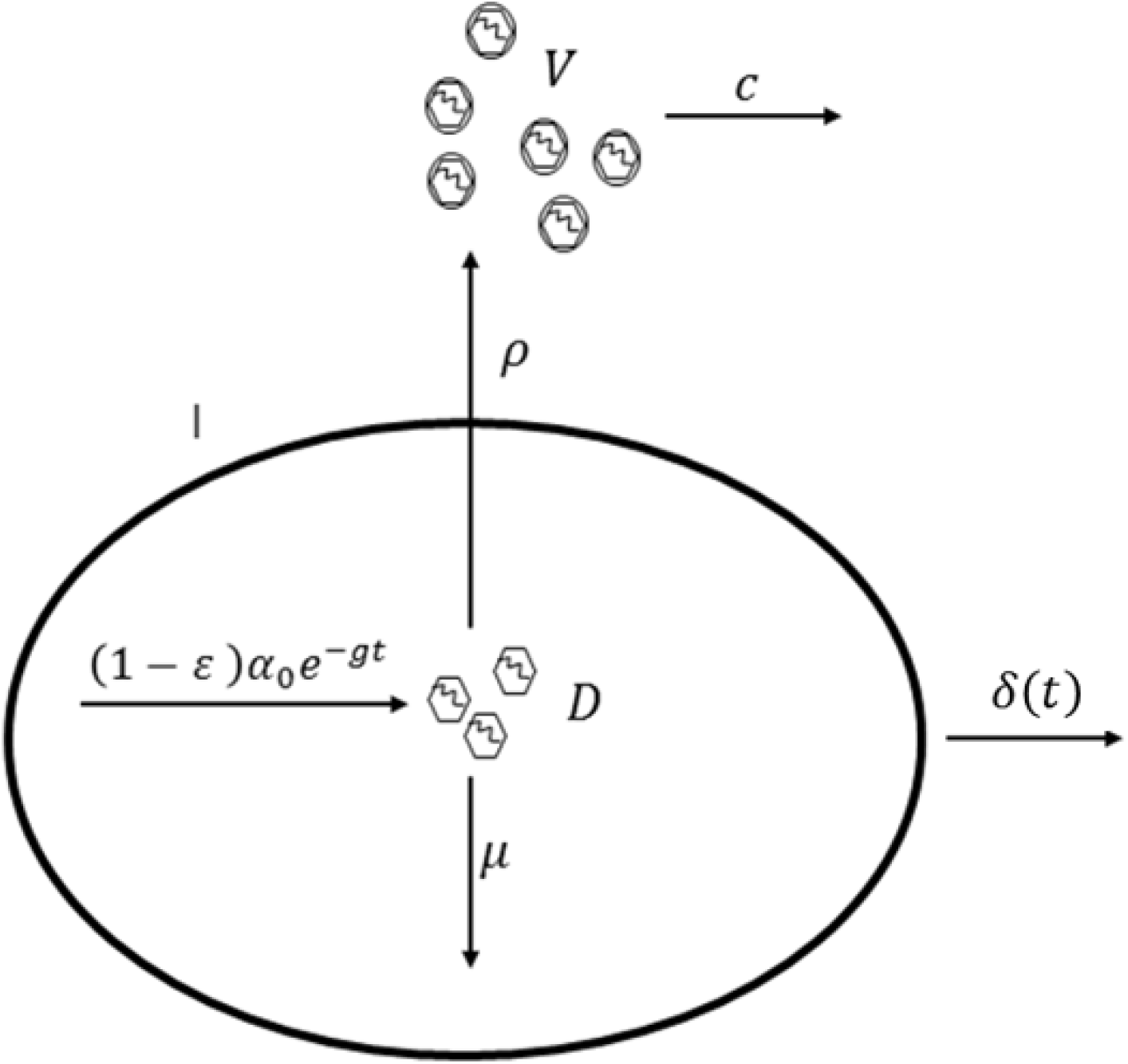
Schematic representation of the HBV multicompartmental model in humanized uPA/SCID chimeric mice. The model includes HBV-infected human hepatocytes (*I*) and die at time-dependent rate *δ*(*t*). Intracellular HBV DNA (*D*) containing capsids are produced with rate constant *α*_0_ and degraded and/or recycled into the nucleus at rate constant *μ*. Capsids are enveloped and secreted using a Hill function 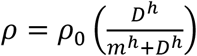. Free virus (*V*) is cleared from blood with rate constant *c*. Antiviral treatment is assumed to reduce the production rate by a fraction (1 – *ε*) and/or assembly/secretion rate by a fraction (1 – *η*) with treatment effectiveness varying between 0 and 1. The exponential decay process, exp(−*gt*), where *g* ≥ 0 is a constant and *t* days post treatment initiation, represents the an additional inhibitory effect on intracellular DNA production.

HBV DNA capsids are enveloped and secreted. Secretion in the model is represented by *ρ*, where 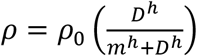. Here *ρ*_0_ is the pre-treatment secretion rate constant, *m* is the *D* level at which the secretion term is 50% of its pre-treatment and *h*, the Hill coefficient is a parameter describing the steepness of the relationship between the decline in *D* during treatment and secretion *ρ* (a justification for including a Hill function instead of using *ρ* as a constant is described in the **Supplementary Information** and (14)).

Infected cells are lost at rate *δ*(*t*) and circulating virus, *V*, is cleared from serum with rate constant *c. B_v_* is the mouse blood volume..

Antiviral treatment can reduce the *D* synthesis rate by 1 – *ε*, slow the viral secretion *ρ* by 1 – *η*, or increase *D* degradation rate by *κ*, where *ε*, *η* and *κ* are parameters that quantify the drug effects (15). Pre-treatment *ε* = 0, *η* = 0 and *κ* = 1 and during antiviral treatment 0 ≤ *ε* ≤ 1, 0 ≤ *η* ≤ land *κ* ≥ 1. We also included an exponential term exp(−*gt*), in the rate of viral synthesis, where *g* ≥ 0 is a constant and *t* the time (days) post treatment initiation, which can account for any additional time-dependent inhibitory effects that may be exerted on intracellular HBV DNA synthesis during treatment, reminiscent of our previous intracellular modelling approach for understanding HCV kinetics under interferon-α treatment (15).

### Parameter estimation and data fitting

To compute the death rate of infected cells, *δ*(*t*), the hAlb decline slope in natural logarithm was calculated. When hAlb levels remained approximately constant an extremely low death rate *δ*(*t*) = *constant* (*δ*(*t*) = 0.001 *d*^−1^) was assumed. In cases where a biphasic decline in hAlb was observed, *δ*(*t*) was estimated using a piecewise function. For instance, in mice with hAlb levels decreasing with rate 0.05 log per day until day 7 from initiation of treatment and that plateaued afterwards, the death rate function was defined as *δ*(*t*) = 0.05 *d*^−1^ for *t* ≤ 7 days and *δ*(*t*) = 0.001 *d*^−1^ otherwise.

We assumed that all human hepatocytes were infected pre-treatment (11, 16, 17) and set *I*_0_ at ~3×10^8^ cells for all mice which was estimated based on each mouse’s approximate hAlb level and weight (not shown). We recently demonstrated that serum and intracellular HBV DNA levels were at steady state for 2 weeks prior to treatment (11). We assume pre-treatment that 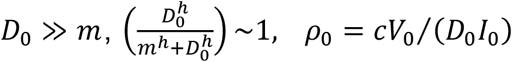 and *α*_0_ = (*ρ*_0_ + *μ*)*D*_0_, where *D*_0_ and *V*_0_ are pre-treatment intracellular and serum HBV DNA levels, respectively. To reduce model uncertainty, we simulated several fixed values of the parameter *μ* as described in Results. We also fixed the volume of distribution to *B_v_* = 1 mL since the blood volume for these relatively small uPA/SCID mice is ~1 mL (18), and fixed the initial intracellular HBV DNA level as *D*_0_=50 copies/cell, approximately the average observed in the mice sacrificed before treatment.

We performed a practical identifiability analysis studying local sensitivity and parameter collinearity. Sensitivity of model output to parameters represents how a change in a parameter value impacts the model output (here *V* and *D* kinetics) (19, 20). Collinearity measures the linear dependence between the parameters. If all the parameters are orthogonal, then the collinearity index equals 1. Alternatively, the collinearity index equals infinity if the parameters are linearly dependent. As a rule of thumb, it is accepted that identifiable models show a collinearity index <20) (19). We used the package FME for the practical identifiability analysis (20).

We used the maximum-likelihood method implemented in MONOLIX version 2016R1 (www.lixoft.com) to estimate the fixed and random effect of the model parameters *c, ε, g, m, h*, and *V*_0_. The fixed effects represent the population average whereas the random effects represent the inter-individual variability.

We estimated the individual parameters as empirical Bayes estimates. Further details about mixed effect models and the population approach used, as well as other details about parameter estimation are given in the **Supplementary Information**. The effect of treatment was tested with a Wald test (21). However, due to the relatively small number of mice per group, this approach can lead to an inflated type I error, therefore we followed Laouénan’s recommendation to compute a corrected p-value (Pc) (**Supplementary Information**).

We compared different models using the Bayesian information criterion (BIC) with the rule “the smaller the better” (22), as well as goodness-of-fits plots. We also considered how accurately the parameters were estimated by analyzing their relative standard error (RSE).

## Results

### Serum and intracellular HBV DNA kinetics

Steady state viral load, *V*_0_, at the onset of treatment ranged between 7.8 and 9.9 log_10_ IU/mL with an average of 9.2 ± 0.4 log_10_ IU/mL (**Figure 2 and Supplementary Table 1**). In mice treated with pegIFN alone, there was a 1-day delay in treatment response after treatment initiation, during which serum HBV DNA remained at pre-treatment level. Then, in all mice a biphasic decline in serum HBV DNA was observed (**Figure 2**), consisting of a short and rapid decrease (1^st^ phase: days 1 - 3 of treatment) followed by a slower decline (2^nd^ phase: days 3 - 14 of treatment). Surprisingly, there was no significant difference in viral kinetics among the treatment groups during the first phase suggesting that initially all antiviral treatments inhibited HBV DNA synthesis to similar levels. In contrast, the log decline in viral load (and the slope) during the 2^nd^ phase was significantly (p = 0.01) different among treatment groups with an average drop of 0.74 ± 0.55 IU/ml (0.06±0.04 IU/day), 0.82 ± 0.37 IU/ml (0.08±0.03 IU/day) and 1.66 ± 0.23 IU/ml (0.15±0.02 IU/day) for pegIFN, LAM and pegIFN+LAM, respectively (**Supplementary Table 1**).

**Figure 2:**
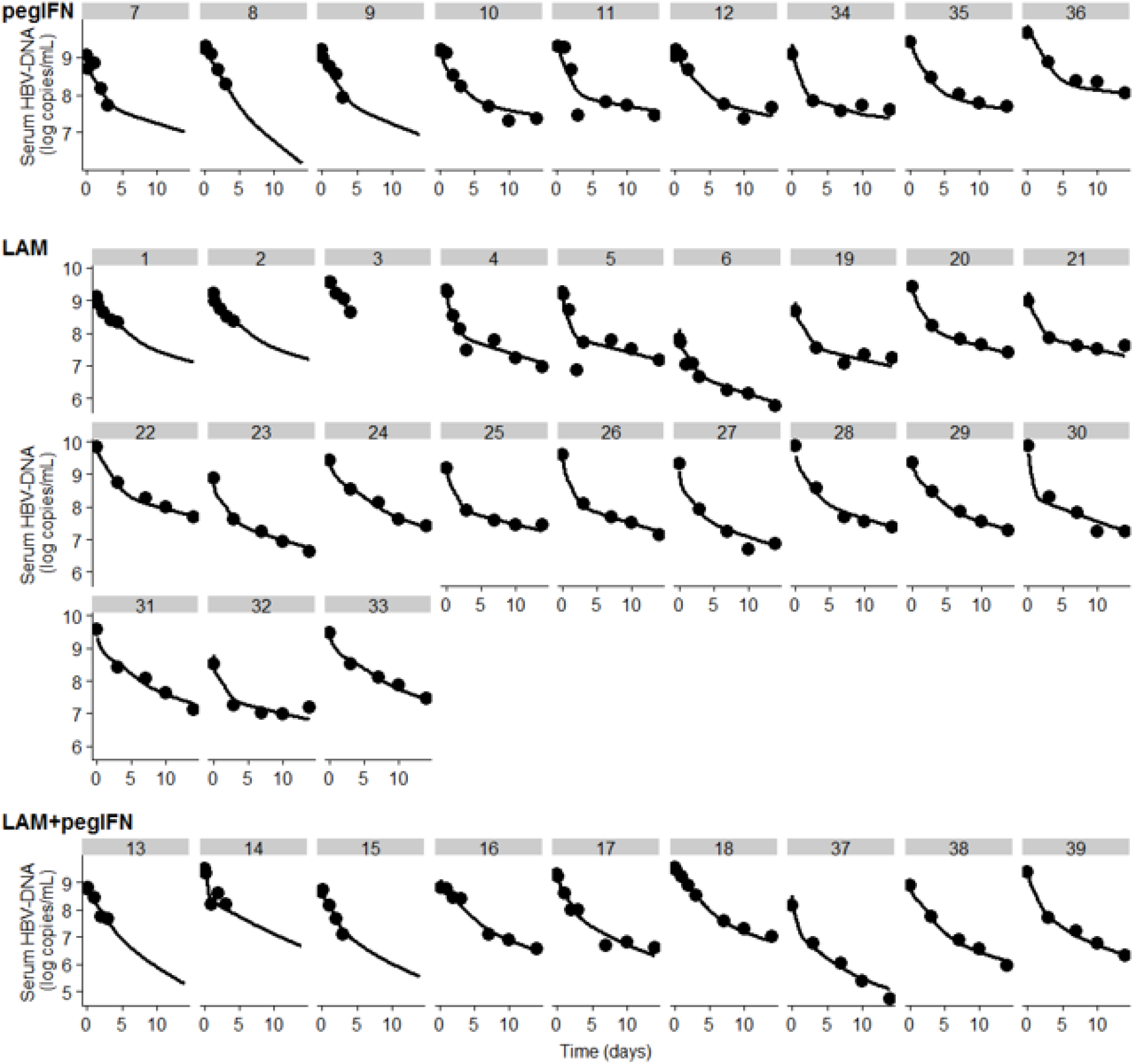
Individual model fit curves (lines) and measured of serum HBV-DNA (dots). Each box represents a mouse. Mice treated with pegIFN are shown in the upper panel, with LAM in the middle panel and with pegIFN+LAM in the lower panel.

The mean level of total intracellular HBV DNA at treatment onset, *D*_0_, was 52 ± 18 copies/cell. At day 3, after the first rapid drop, the average intracellular HBV DNA was 36, 26 and 38 copies/cell in the pegIFN, LAM and the pegIFN+LAM group, respectively (see Methods). After the second slower drop, the average intracellular HBV DNA at day 14 was 1, 17 and 9 copies/cell under pegIFN, LAM and pegIFN+LAM, respectively (**Figure 3**).

**Figure 3:**
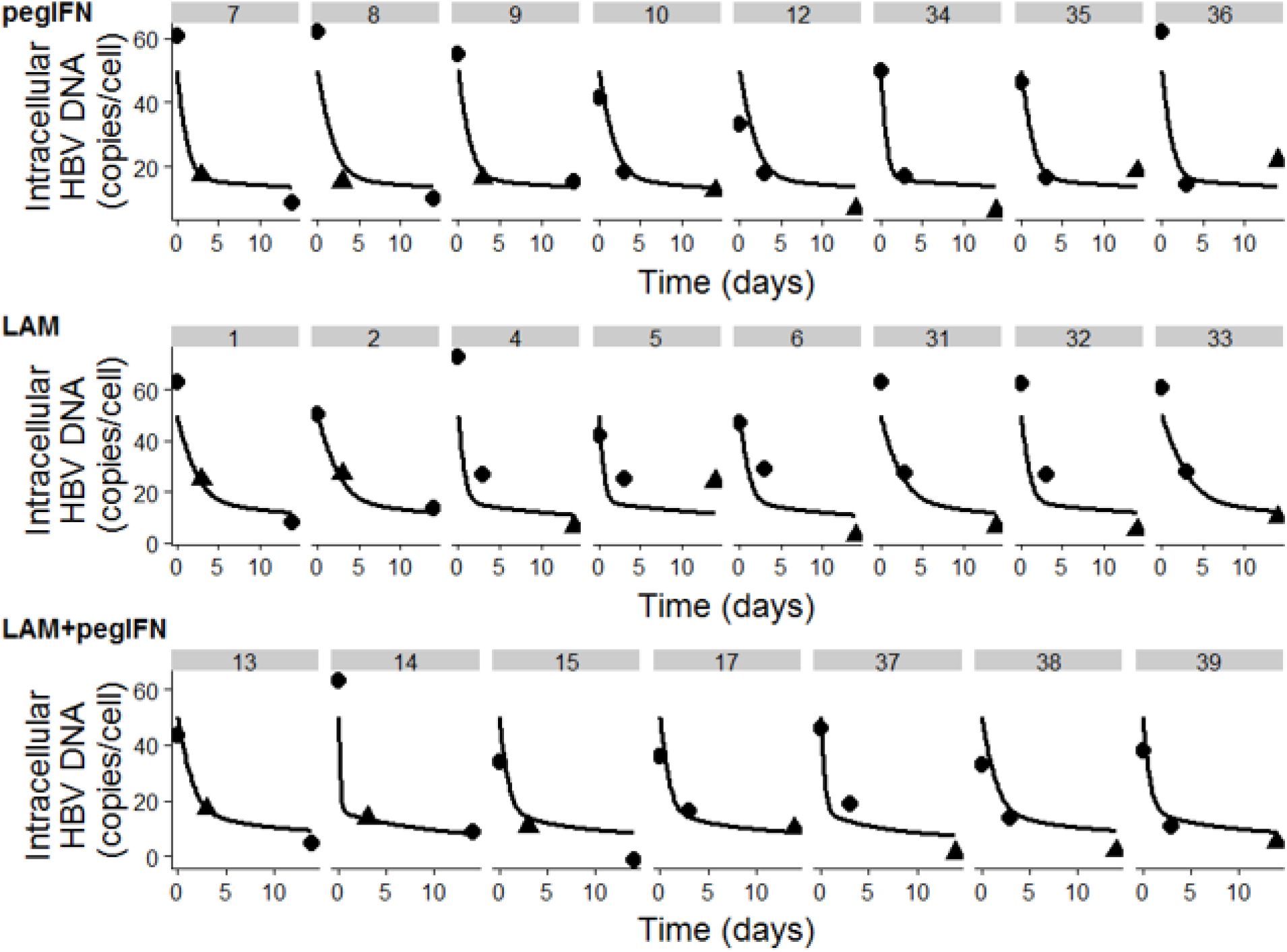
Individual model fit curves (lines) and measured of intracellular HBV-DNA. Each box represents a mouse. Mice treated with pegIFN are shown in the upper panel, with LAM in the middle panel and with pegIFN+LAM in the lower panel. Observed intracellular HBV-DNA is shown by a triangle and simulated data by a dot.

### Serum human albumin kinetics

Pre-treatment the hAlb level in the serum was 8.3 ± 2.6 mg/mL on average with no significant difference among treatment groups (**Supplementary Table 2**). During LAM administration hAlb levels remained stable in the serum of all mice (**Supplementary Figure 1** and **Supplementary Table 2**) with an average kinetic slope of 0.02 ± 0.19 mg/mL/d which is not significantly different from a plateau (i.e., slope = 0). In contrast, mice treated with pegIFN or pegIFN±LAM showed an initial decline in hAlb followed by a plateau or slight increase/decrease (**Supplementary Figure 1**). The first decline phase persisted 7-10 days after treatment initiation and the average decrease rate was 0.68 ± 0.56 and 0.61 ± 0.61 mg/mL/d, under pegIFN or pegIFN±LAM, respectively (**Supplementary Table 2**). During the second phase, the median rate of change in hAlb was 0.01 (p=0.20, range, −0.27;0.22) mg/mL/d (**Supplementary Table 2**).

### The main MOA of PegIFN is blocking HBV DNA synthesis

To predict pegIFN MOA and estimate viral-host parameters under pegIFN, LAM and pegIFN+LAM we next performed model (**Equation1**) sensitivity analysis against both the intracellular and extracellular (serum) measured data. We show that the model does not reproduce the expected effect of LAM in blocking HBV DNA synthesis solely by using parameter *ε* or parameter *g* alone (**Supplementary Figure 2**), but only when both are assumed to be inhibited (**Supplementary Figure 3**).

We then used the model to test different possible MOAs for pegIFN. We simulated models where pegIFN: (i) blocks virus secretion, *η;* (ii) blocks HBV DNA synthesis, *ε;* or (iii) enhances the HBV DNA degradation rate as *κμ*. However, none of these MOA separately captured the measured viral kinetics of both intracellular and serum HBV DNA. Indeed, the model with inhibition of virus secretion led to an increased intracellular HBV DNA and to a rebound in serum HBV DNA that are inconsistent with our observations. Models with *ε* >0 or *κ* >1 did not reproduce the data (**Supplementary Figure 4**). However, similar to the LAM data, best fits were achieved when pegIFN (with and without LAM; **Supplementary Figure 5**) was simulated to act mainly by blocking HBV DNA synthesis (*ε*) with an additional time-dependent inhibition (parameter *g*), which represents a time-dependent inhibition of viral DNA synthesis. As such, in the following model investigations we set *κ* = 1 and *η* = 0.

### Establishing a biologically plausible intracellular HBV DNA degradation rate constant

Model sensitivity analysis results of the remaining unknown parameters μ, *c, m, h, g* and *ε* suggest that collinearity index was 13 (i.e., the model is identifable) only for models where μ was fixed (**Supplementary Figure 6**). We found that an intracellular DNAs clearance rate, 0.01 ≤ *μ* ≤ 0.05 *d*^−1^ (corresponding of intracellular HBV DNA *t*_1/2_ between 14 hr and 69 hr which are in agreement with Xu *et al*. estimate of *t*_1/2_~24 *hr* (23)) provide the most plausible estimates of serum HBV DNA *t*_1/2_ (see **Sensitivity Analysis** in **Supplementary Information**). To be conservative, we set *μ* = 0.01 *d*^−1^ (the longest intracellular DNAs half-life) for the following modelling sections.

### Serum and intracellular viral-host parameter estimations

Importantly, simultaneously fitting the model to both serum and intracellular HBV DNA kinetic data yielded good agreement between the best-fit model curve and the data (**Figures 2 & 3**) and the goodness of fit diagnostic was satisfactory (not shown) allowing us to estimate key viral-host parameters (**Table 1**). The initial serum HBV DNA viral load, *V*_0_ was estimated as 9.3 log_10_ copies/mL (RSE=0.7%). The serum virus clearance rate, *c*, was estimated as 6.13 d^−1^ (RSE=24%), leading to an average serum HBV DNA half-life of 2.7 hr.

**Table 1:**
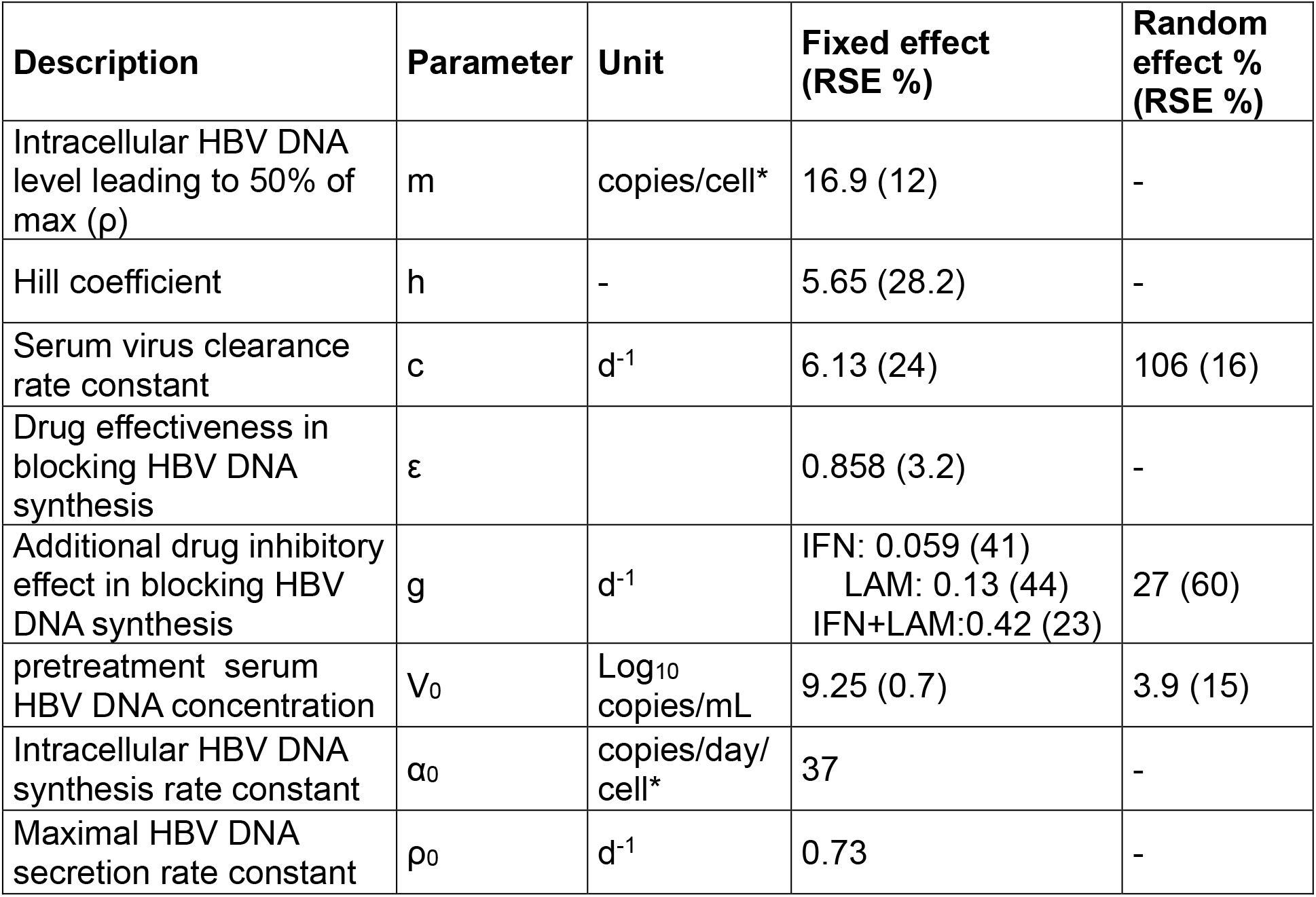
Model (Equation 1) parameter estimateations. Fixed effects represents the average population value and random effect the inter-individual variability (IIV). RSE stands for relative standard error. Additive error for V: 0.19 log_10_ copies/mL. Additive error for D: 6.08 copies/cell. Intracellular HBV DNA degaradation rate constant μ was set to 0.01 d^−1^, the initial intracellular HBV DNA was set to 50 copies/cell and the volume of distrbution to 1 mL. * assuming that the initial number of infected cells, *I*_0_=3×10^8^ cells.

Intracellular HBV DNA synthesis rate constant *α*_0_ and pre-treatment virion excretion rate constant *ρ*_0_ were estimated to be 37 copies/day/cell and 0.73 d^−1^, respectively (**Figure 4A**). We also estimated the parameters describing the relationship between *p* and intracellular HBV DNA levels as *m* = 16.9 copies/cell (RSE=12%) and h=5.7 (RSE=28%). This implies that during treatment virus secretion rate is 50% of its pre-treatment value (i.e., steady state) when the intracellular HBV DNA level drops to 17 copies/cell (**Figure 4B**).

**Figure 4:**
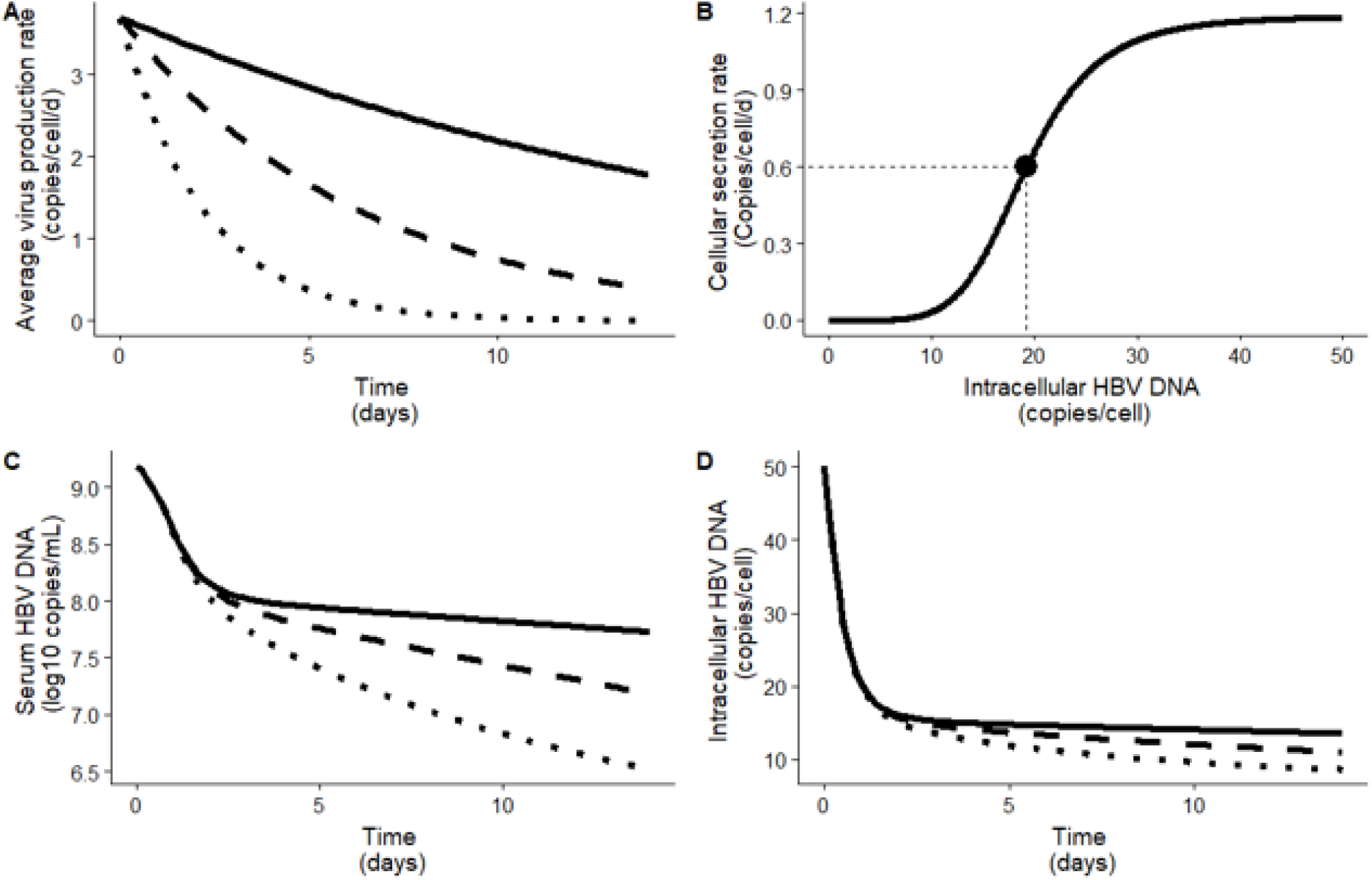
Predicted dynamic of HBV infection in chimeric mice. The predictions were simulated using the population parameters shown in **Table 1** for the different treatments: pegIFN (solid line), LAM (dashed line) and PegIFN+LAM (dotted line). (**A**) Average intracellular HBV DNA production rate under treatment, computed as (1 – *ε*)*α*_0_*e*^−*gt*^. (**B**) Predicted virus secretion rate, i.e. 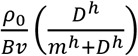, according to intracellular HBV DNA, D, decline from pre-treatment level *due to treatment*. The dot represents the inflection point where the secretion rate is divided by 2 when intracellular level of HBV DNA is m=16.9 copies/cell. (**C**) Predicted average dynamics of serum HBV DNA, *V*. (**D**) Predicted average dynamics of intracellular HBV DNA, *D*.

The average treatment effectiveness in blocking HBV DNA synthesis, *ε*, for all treatments was estimated as 85.8 % (RSE=3.2%). However, the additional treatment inhibitory effect on intracellular HBV DNA synthesis, *g*, was significantly different between the treatment groups with *g* = 0.059 *d*^−1^ for pegIFN, 0.13 d^−1^ for LAM (Pc=0.026) and 0.42 d^−1^ for pegIFN+LAM (Pc<0.001) (**Table 1**). This translates into a faster decline of intracellular HBV DNA and consequently of serum HBV DNA for mice treated with pegIFN+LAM compared to the two other treatment groups (**Figures 4 C & D**). Lastly, there was no association between donor human hepatocytes IL28B genotypes and estimated HBV kinetic parameters (not shown).

## Discussion

The typical HBV DNA decline in serum of patients treated with pegIFN and/or LAM is biphasic, which consists of a first rapid decline phase followed by a slower 2^nd^ phase (6–8). It was hypothesized that the first phase represents treatment efficacy in blocking HBV DNA production and serum virus half-life while the slower second phase represents loss/death of infected cells. However, in the current study, a biphasic HBV decline was also observed in serum of humanized mice treated with pegIFN and/or LAM in the absence of adaptive immune response. Interestingly, we observed that while the first phase was similar in time and magnitude under pegIFN and/or LAM, the 2^nd^ decline phase exhibited significantly different reductions in serum HBV DNA level among the three treatment regimens with pegIFN, LAM and pegIFN+LAM showing an increasing order of antiviral potency. While the intracellular HBV DNA appears to show a similar biphasic decline, suggesting that the observed 2^nd^ phase decline in patients may reflect intracellular HBV DNA loss rather than death of infected hepatocytes, more frequent sampling will be needed to confirm such a conclusion and assess the slopes of the different phases more precisely. Regardless, the kinetic picture obtained herein in both serum and within human hepatocytes provides a unique opportunity to investigate HBV dynamics that is not feasible in treated patients.

To provide insights into the dynamics of HBV treatment response and the mode of action of interferon-α against HBV, we developed a multicompartmental model to describe both the extracellular (serum) and intracellular HBV DNA kinetic data. The model predicts that pegIFN mainly inhibits intracellular HBV DNA synthesis in humanized uPA/SCID mice with an efficacy of 86%. While the initial efficacy of blocking HBV DNA synthesis was similar under pegIFN, LAM and pegIFN+LAM, the model suggests that pegIFN inhibition of intracellular HBV DNA synthesis is associated with a slower secondary inhibitory effect than LAM, leading to slower second phase decline. However, the molecular basis of the slower secondary inhibitory effect remains to be determined.

We also monitored the degree of functional human hepatocyte repopulation by measuring hAlb in mouse serum. During LAM therapy, hAlb levels remained constant. However, a biphasic decrease in hAlb occurred during regimens that included pegIFN. This hAlb kinetics was also observed in HCV infected chimeric mice undergoing pegIFN treatment (unpublished data). Therefore, the reduction is independent of HBV infection. Because some ISGs can lead to apoptosis of virus-infected cells (24–27) and many ISGs have been shown to be induced by interferon −α treatment in the human hepatocytes of the chimeric mice (28), one possibility is that the reduction of hAlb levels by pegIFN could be due to human hepatocyte apoptosis. However, an alternative explanation for the decline in hAlb under pegIFN could be changes in gene expression that are unrelated to cell death. To account for these two possibilities, we first modeled infected cell loss as a function of hAlb decline and then modified the Eq. 1 assuming no enhanced death/loss by pegIFN (i.e., ignoring hAlb kinetics), and calibrated it with observed HBV DNA kinetics. Importantly, we found that the main model results do not significantly change (see Supplementary Information), although the modified model has additional unknown parameter to estimate. Further experimental analysis (e.g. ALT measurements, gene expression studies) are needed to reveal the exact mechanism by which interferon causes a decline in serum hAlb in humanized mice.

Modeling results predict a plausible circulating serum virus half-life in chronic HBV-infected mice with humanized liver of ~2.7 hr, consistent with our previous estimate of virus half-life of ~1 hr during acute infection in the same humanized mouse model (11). Our estimate is also comparable with previous virion half-lives estimated in chimeric mice which are 2.6 hr in high viremic uPA mice and 67 min in low viremic uPA mice (29). However, our serum virus half-life estimate of 2.7 hr in HBeAg+ humanized mice is significantly shorter than in HBeAg+ humans treated with LAM for whom virus half-life was estimated to be 17 hours (30) or 1 day (31). As we have previously demonstrated for hepatitis C virus (HCV) (32), the calculation of viral serum half-life is much more precise using drugs that specifically and rapidly block viral secretion (15, 33). Hence, in the future, this discrepancy may be better investigated/resolved using new HBV antivirals that block HBV secretion.

Murray et al. developed within-host dynamics model for HBV using serum and liver samples collected during acute HBV infection from chimpanzees (34). In the current study, we model the serum and intracellular HBV kinetics under antiviral treatment in chronically HBV-infected humanized mice and provide novel insights into intracellular HBV dynamics. Assuming that ~3×10^8^ human hepatocytes were productively HBV-infected pre-treatment, the model predicts that at steady state the intracellular HBV DNA synthesis rate is ~37 copies per cell per day and that ~27 HBV DNA copies (i.e. DNA-containing capsids) are secreted per cell per day. That being said, one limitation of the current study was the infrequent intracellular HBV kinetic data. In this model intracellular HBV DNA was only measured once at the time each mouse was euthanized. Thus, we only observed one value per animal at either day 0, 3 or 14. Although we imputed the two missing values for each mouse using uniform distribution based on the intracellular HBV DNA levels observed in the other mice from the same treatment group, this still provides relatively infrequent sampling, a limitation that could possibly be addressed in future experiments. In addition, the limitation to draw large blood volume in the current humanized mouse model precluded kinetic data of drug pharmacokinetics, HBsAg and ALT.

In summary, we developed a multicompartmental mathematical model to describe both the serum and intracellular HBV treatment response kinetics in the humanized uPA/SCID mice as a means to reveal drug mechanism of action and key viral-host parameters. Analogous to LAM, our model predicts that pegIFN mainly inhibits HBV DNA synthesis in the humanized uPA/SCID mice and that the combination of pegIFN+LAM induces higher secondary time-dependent inhibition of HBV DNA synthesis compared to monotherapy with LAM or pegIFN. Reminiscent of observations in patients treated with pegIFN and/or LAM, a biphasic HBV decline was observed in treated humanized mice in the absence of adaptive immune response, suggesting that the observed 2^nd^ phase decline in patients may reflect intracellular HBV DNA loss rather than death of infected hepatocytes. Notably, these steady state perturbations allow us to make novel estimates of intracellular HBV dynamics in human hepatocytes suggesting that ~37 HBV DNA copies are synthesized per cell per day of which 73% are secreted. The model further suggests that secretion rate is coupled to intracellular HBV DNA, with best fit being achieved using a Hill function. How these dynamics change during treatment would be interesting to determine and perhaps help reveal the nature of the secondary time-dependent inhibition which distinguishes the treatment regimens analyzed in this study.

