## Supplementary File for "Understanding hepatitis B virus dynamics and the antiviral effect of interferon-α treatment in humanized chimeric mice"

(1) The Program for Experimental and Theoretical Modeling, Division of Hepatology, Department of Medicine, Loyola University Medical Center, Maywood, IL, USA; (2) Centre for Immunity, Infection and Evolution, University of Edinburgh, UK; (3) Liver Research Project Center, Hiroshima University, Hiroshima, Japan;(4) PhoenixBio Co., Ltd., Hiroshima, Japan; (5) Department of Gastroenterology and Metabolism, Graduate School of Biomedical & Health Sciences, Hiroshima University, Hiroshima, Japan; (6) Department of Microbiology and Immunology, Loyola University Medical Center, Maywood, IL, USA; (7) Institut für Biostatistik and Mathematische Modellierung, Fachbereich Medizin, Goethe Universität, Frankfurt, Germany; (8) Theoretical Biology and Biophysics, Los Alamos National Laboratory, Los Alamos, NM, USA

*These authors contributed equally to this study

[#]corresponding authors

Kazuaki Chayama,
Department of Gastroenterology and Metabolism, Institute of Biomedical and Health Sciences,
Hiroshima University 1-2-3, Kasumi, Minami-ku, Hiroshima-shi, Hiroshima,734-8551, Japan
P:+81-82-257-5190
F:+81-82-255-6220


Harel Dahari,
Program for Experimental and Theoretical Modeling, Division of Hepatology, Department of Medicine, Loyola University Medical Center, Maywood, IL, USA.
P: +1-708-216-4682
F: +1-708-216-6299
;

**A. Model selection**

1. Modeling the role hAlb as surrogate marker of HBV-infected cells death.

In the model (Eq. 1) we assumed a time-dependent $\delta(t)$ as a function of hAlb slopes assuming that a decline in hAlb under pegIFN reflects HBV-infected cell death. However, the decline in hAlb could equally be due to pegIFN-induced changes in gene expression without cell death. To explore this scenario, we assumed in Eq. 1 that $\delta(t) = constant\ \delta$. The modified model was calibrated with fixed parameters and unknown parameters as done in Figs 2 and 3 in the main text. The BIC were similar for Eq. 1 and modified Eq. 1 models (i.e. 694 and 691, respectively). However, the model (Eq. 1) with $\delta(t)$ as a function of hAlb slopes requires one parameter less to estimate compared to a model with constant $\delta$ (which was estimated $\delta = 0.0808\ d^{-1}$) and was therefore chosen as the model used in our study.

2. Modeling HBV DNA secretion rate

We fit two models, with and without the Hill function, to describe the HBV DNA secretion rate constant and found BIC of 709 and 771, respectively. As this is also consistent with our recent findings in primary human hepatocytes (1) we therefore ruled out the model without the Hill function.

**B. Sensitivity analysis**

Because of the non-sensitivity of the model (Eq. 1) to $\mu$, we tested different models where we fixed the HBV DNA degradation $\mu$ to different values between 0.001 and 10 $d^{-1}$ (**Supplementary Figure 7**). The parameters $V_0$, $c$, $m$, $h$, $g$ and $\varepsilon$ were free and estimated. We identified three intervals: A) $\mu < 0.075\ d^{-1}$, B) $\mu \in [0.075 - 1.00]$ $d^{-1}$ and C) $\mu > 1.00$ $d^{-1}$. For $\mu \in [0.075 - 1.00]$ $d^{-1,}$ the parameters $g$, $m$ and $h$ were poorly estimated with relative standard error > 40% and the model did not converge for $\mu$ between 0.3 and 0.4 $d^{-1}$. The initial serum viral load, $V_0$, proved to be robust against changes in $\mu$ and varied between 9.13 for $\mu = 0.5\ d^{-1}$ and 9.26 for $\mu = 0.001\ d^{-1}$. In the intervals A and C, the additional time-dependent inhibitory effect under pegIFN+LAM and LAM, modeled by the parameter $g$, were significantly different from the estimated value of $g$ for pegIFN alone. The intracellular DNA level leading to 50% of maximal secretion rate, $m$, proved to be robust against changes in $\mu$ in A and C, varying between 16.5 and 18 copies/cell and between 53.2 and 57.5 copies/cell, respectively. The serum HBV DNA clearance rate, $c$, the Hill factor, $h$, the drug effectiveness in blocking HBV DNA synthesis, $\varepsilon$, and the additional time-dependent inhibitory effect on HBV DNA synthesis, $g$, under pegIFN+LAM treatment decreased with increasing $\mu$. This leads the serum HBV DNA free half-life $(t_{1/2})$ to vary between $t_{1/2} =$

$1.1\ hr$ for $\mu = 0.01\ d^{-1}$. and $t_{1/2} = 14.7\ hr$ for $\mu = 10\ d^{-1}$. Interestingly, the shorter serum HBV $t_{1/2}$ of 1.1 hr is in agreement with our recent estimate ($t_{1/2} \sim 1\ hr$) during acute infection (2). Concerning the Hill factor, $h$, the drug effectiveness, $\varepsilon$, and free virus clearance, $c$, higher values are observed in interval A than in interval C (**Supplementary Figure 7**). More specifically, $h$ varied between 4.36 and 6.60 in A and between 0.51 and 1.03 in C; $\varepsilon$ varies between 0.828 and 0.895 in A and between 0.546 a,d 0.571 in C; $c$ varies between 4.73 and 7.17 $d^{-1}$ (corresponding to free virus half-life varying between 2.33 and 3.51 hr) in A and between 1.11 and 1.21 $d^{-1}$ (corresponding to free virus half-life varying between 13.8 and 14.9 hr) in C.

### Supplementary Table 1: Virus kinetic analysis

| Treatment | Days of treatment | Donor | ID | Intracellular HBV DNA measured | V0 [log IU/mL] | Drop in VL [log10] |
|---|---|---|---|---|---|---|
| LAM | 14 | 2YM | 23 | N | 8.88 | -1.33 |
| LAM | 14 | 2YM | 24 | N | 9.41 | -1.08 |
| LAM | 14 | 2YM | 25 | N | 9.19 | -1.57 |
| LAM | 14 | 1YF | 19 | N | 8.67 | -1.58 |
| LAM | 14 | 1YF | 20 | N | 9.43 | -1.41 |
| LAM | 14 | 1YF | 21 | N | 8.99 | -1.41 |
| LAM | 14 | 1YF | 22 | N | 9.82 | -1.26 |
| LAM | 14 | 5YM | 31 | Y | 9.25 | -2.25 |
| LAM | 14 | 5YM | 32 | Y | 8.5 | -1.62 |
| LAM | 14 | 5YM | 33 | Y | 9.18 | -1.86 |
| LAM | 14 | 5YM | 26 | N | 9.6 | -1.59 |
| LAM | 14 | 5YM | 27 | N | 9.32 | -2.11 |
| LAM | 14 | 5YM | 28 | N | 9.87 | -2.11 |
| LAM | 14 | 5YM | 29 | N | 9.37 | -1.34 |
| LAM | 14 | 5YM | 30 | N | 9.86 | -1.92 |
| LAM | 3 | 2YF | 1 | Y | 8.97 | -3.50 |
| LAM | 3 | 2YF | 2 | Y | 9.07 | -3.64 |
| LAM | 3 | 2YF | 3 | N | 9.56 | -4.14 |
| LAM | 14 | 2YF | 4 | Y | 9.27 | -1.60 |
| LAM | 14 | 2YF | 5 | Y | 9.34 | -1.87 |
| LAM | 14 | 2YF | 6 | Y | 7.83 | -0.85 |
| **LAM Mean (SD)** | | | | | **9.21 (0.50)** | **-1.91 (0.85)** |
| pegIFN | 14 | 5YM | 34 | Y | 9.08 | -1.48 |
| pegIFN | 14 | 5YM | 35 | Y | 9.40 | -1.30 |
| pegIFN | 14 | 5YM | 36 | Y | 9.65 | -1.12 |
| pegIFN | 3 | 2YF | 7 | Y | 8.97 | -5.59 |
| pegIFN | 3 | 2YF | 8 | Y | 9.31 | -4.61 |
| pegIFN | 3 | 2YF | 9 | Y | 9.14 | -5.21 |
| pegIFN | 14 | 2YF | 10 | Y | 9.25 | -1.56 |
| pegIFN | 14 | 2YF | 11 | N | 9.62 | -1.85 |
| pegIFN | 14 | 2YF | 12 | Y | 9.19 | -1.82 |
| **pegIFN mean (SD)** | | | | | **9.29 (0.23)** | **-2.72 (1.84)** |
| pegIFN+LAM | 14 | 5YM | 37 | Y | 7.80 | -3.28 |
| pegIFN+LAM | 14 | 5YM | 38 | Y | 8.90 | -1.63 |
| pegIFN+LAM | 14 | 5YM | 39 | Y | 8.83 | -2.80 |
| pegIFN+LAM | 3 | 2YF | 13 | Y | 8.78 | -5.74 |
| pegIFN+LAM | 3 | 2YF | 14 | Y | 9.23 | -5.46 |
| pegIFN+LAM | 3 | 2YF | 15 | Y | 8.70 | -7.46 |
| pegIFN+LAM | 14 | 2YF | 16 | N | 8.97 | -1.83 |
| pegIFN+LAM | 14 | 2YF | 17 | Y | 9.19 | -2.42 |
| pegIFN+LAM | 14 | 2YF | 18 | N | 9.51 | -1.84 |
| **pegIFN**+LAM **mean (SD)** | | | | | **8.88 (0.48)** | **-3.61 (2.10)** |
| **Overall mean (SD)** | | | | | **9.15 (0.45)** | **-2.49 (1.59)** |

**Supplementary Table 2: Serum human albumin kinetic parameters**

| Subject | Donor | Treatment | Slope s1 (mg.ml$^{-1}$.d$^{-1}$) | End of phase 1 (d) | Slope s2 (mg.ml$^{-1}$.d$^{-1}$) |
|---|---|---|---|---|---|
| d2m102 | 2YM | LAM | 0.03 | NA | NA |
| d2m103 | 2YM | LAM | -0.04 | NA | NA |
| d2m104 | 2YM | LAM | 0.09 | NA | NA |
| d5m101 | 5YM | LAM | -0.06 | NA | NA |
| d5m103 | 5YM | LAM | -0.04 | NA | NA |
| d5m104 | 5YM | LAM | -0.04 | NA | NA |
| d5m202 | 5YM | LAM | -0.12 | NA | NA |
| d5m203 | 5YM | LAM | -0.10 | NA | NA |
| d1m101 | 1YF | LAM | -0.08 | NA | NA |
| d1m102 | 1YF | LAM | -0.05 | NA | NA |
| d1m103 | 1YF | LAM | -0.04 | NA | NA |
| d1m104 | 1YF | LAM | 0.02 | NA | NA |
| d5m23 | 5YM | LAM | 0.02 | NA | NA |
| d5m24 | 5YM | LAM | -0.03 | NA | NA |
| d5m25 | 5YM | LAM | 0.06 | NA | NA |
| 101 | 2YF | LAM | 0.27 | NA | NA |
| 102 | 2YF | LAM | 0.74 | NA | NA |
| 103 | 2YF | LAM | -0.30 | NA | NA |
| 104 | 2YF | LAM | 0.07 | NA | NA |
| 105 | 2YF | LAM | -0.03 | NA | NA |
| 106 | 2YF | LAM | -0.05 | NA | NA |
| d5m26 | 5YM | pegIFN | -0.35 | 10 | 0.02 |
| d5m27 | 5YM | pegIFN | -0.39 | 10 | 0.22 |
| d5m28 | 5YM | pegIFN | -0.48 | 7 | 0.00 |
| 201 | 2YF | pegIFN | -0.29 | 3 | NA |
| 202 | 2YF | pegIFN | -2.07 | 3 | NA |
| 203 | 2YF | pegIFN | -0.68 | 3 | NA |
| 204 | 2YF | pegIFN | -0.30 | 3 | 0.02 |
| 205 | 2YF | pegIFN | -0.84 | 3 | -0.07 |
| 206 | 2YF | pegIFN | -0.68 | 3 | -0.27 |
| d5m29 | 5YM | PegIFN+LAM | -0.10 | 10 | 0.00 |
| d5m30 | 5YM | PegIFN+LAM | -0.66 | 7 | -0.03 |
| d5m31 | 5YM | PegIFN+LAM | -0.47 | 7 | 0.07 |
| 301 | 2YF | PegIFN+LAM | -1.20 | 3 | NA |
| 302 | 2YF | PegIFN+LAM | 0.73 | 3 | NA |
| 303 | 2YF | PegIFN+LAM | -0.42 | 3 | NA |
| 304 | 2YF | PegIFN+LAM | -1.19 | 3 | 0.07 |
| 305 | 2YF | PegIFN+LAM | -0.03 | 7 | -0.13 |
| 306 | 2YF | PegIFN+LAM | -1.60 | 3 | 0.08 |

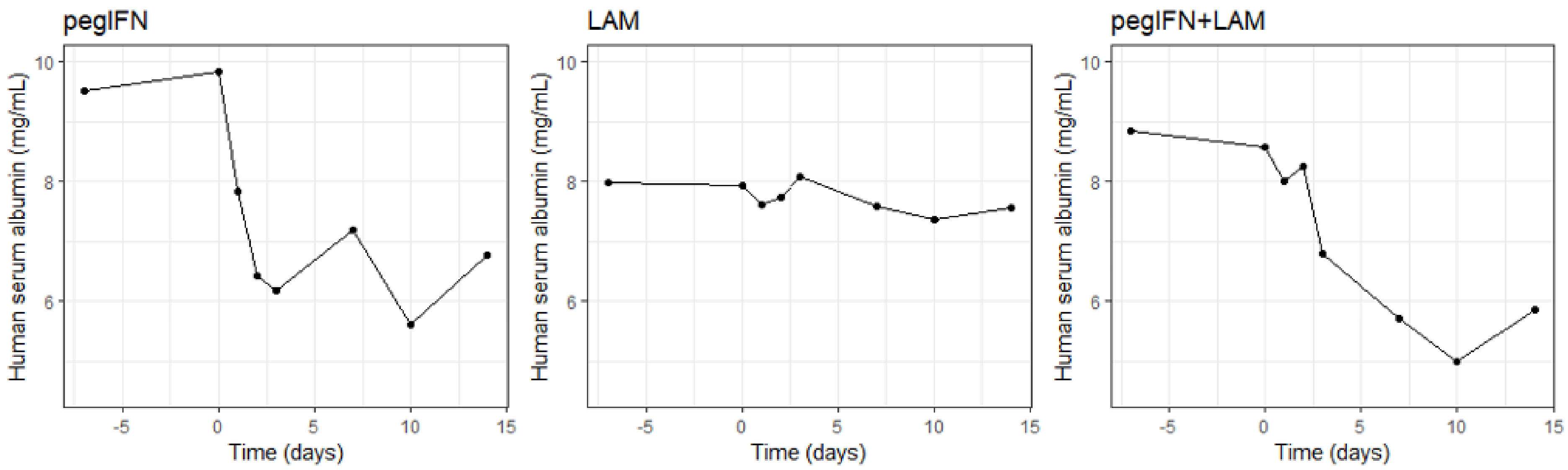


**Supplementary Figure 1. Average serum human albumin kinetics for mice receiving pegIFN monotherapy (left panel), LAM monotherapy (center panel) or pegIFN+LAM combination (right panel).**

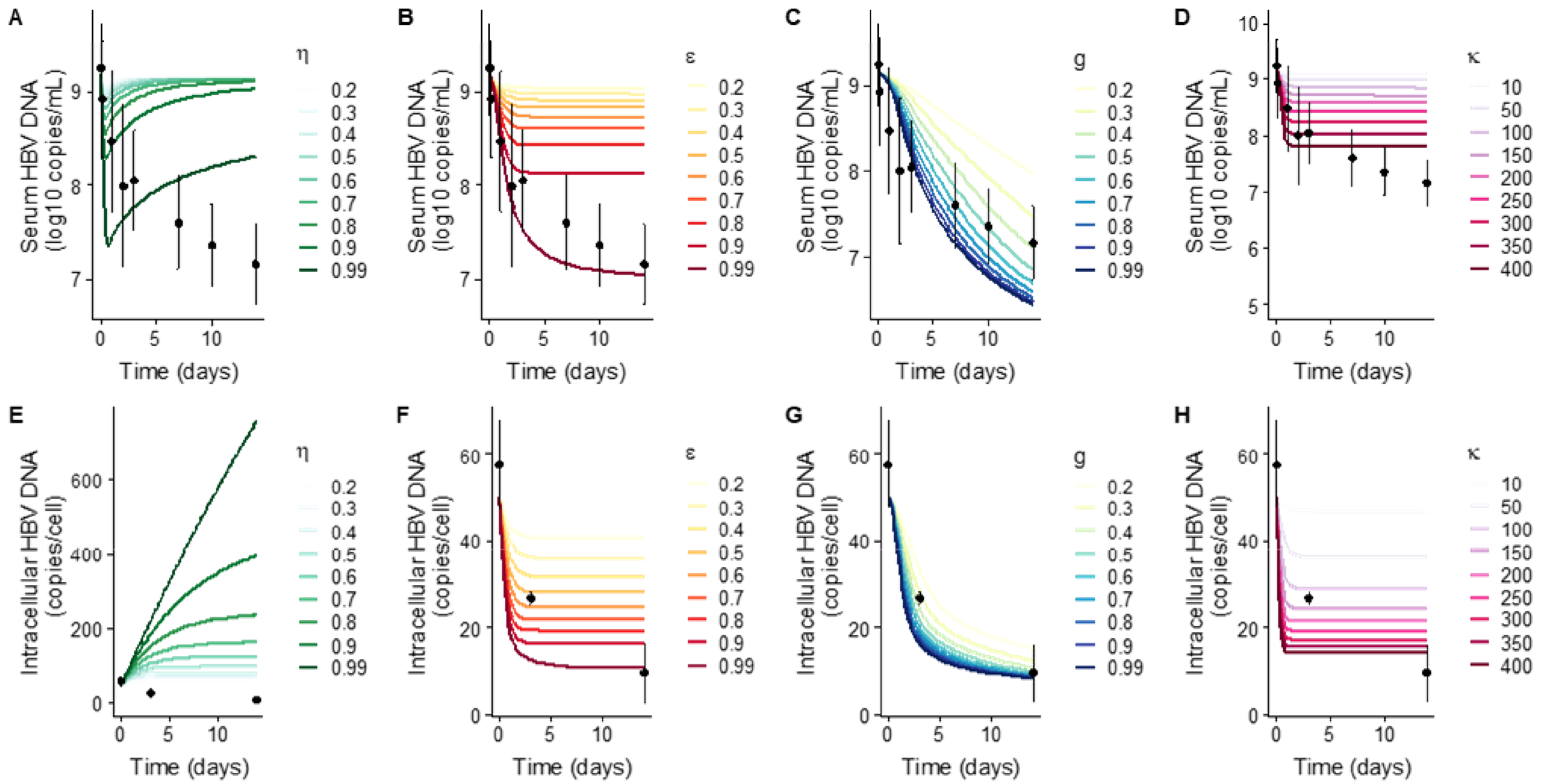


**Supplementary Figure 2: Simulation of serum (A, B, C, D) and intracellular (E, F, G, H) HBV DNA kinetics in LAM group and comparison with observed data using Eq. 1.**

**A, E**: We modelled the treatment effect as the inhibition of secretion as follow: $(1-\eta)\rho$ where η is the drug effectiveness (varying between 0 and 1) and ρ is the secretion rate. g, ε and κ are fixed to 0. **B, F:** We modelled the treatment effect as the intracellular HBV DNA production as follow: $(1-\varepsilon)\alpha_0$ where ε is the drug effectiveness (varying between 0 and 1) and $\alpha_0$ is the intracellular HBV DNA production rate. g, κ and η are fixed to 0. **C, G:** We modelled the additional inhibitory effect on the intracellular HBV DNA production as follow: $\alpha_0 \exp(-g*t)$ where g is the additional inhibitory effect and $\alpha_0$ is the intracellular HBV DNA production rate. ε, η and κ are fixed to 0. **D, H**: We modelled the effect of treatment on the degradation rate μ as follow: μ.κ where $\mu$=0.01 $d^{-1}$ and ε and η are fixed to 0. In all simulations the model's parameters were fixed at $I_0$=3.10$^8$ cells, $V_{,0}$= 9.18 $\log_{10}$ copies/mL, $D_{,0}$=50 copies/cell, δ=0.01 $d^{-1}$, $B_v$ =1.0 mL, c=11.8 $d^{-1}$, m=19.2 copies/cell, h=5.47, g=0 $d^{-1}$ and $\mu$=0.01 $d^{-1}$.

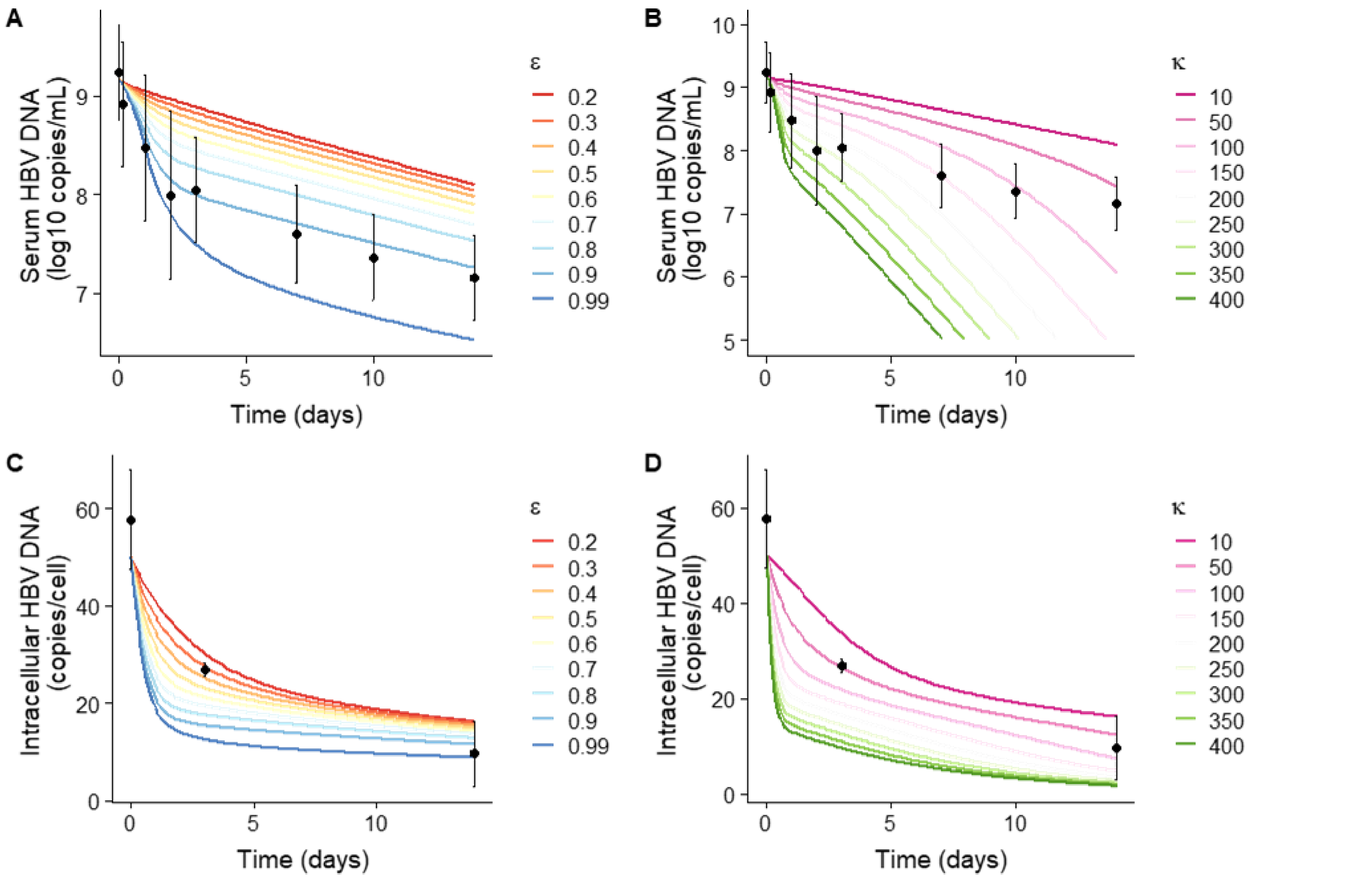


**Supplementary Figure 3. Testing multiple MOA for LAM group. Simulation of serum (A, B) and intracellular (C, D) HBV DNA kinetics in LAM group and comparison with observed data.**

A, C: We modelled the treatment effect as the intracellular HBV DNA production as follow: $(1-\varepsilon)\alpha_0$ where ε is the drug effectiveness (varying between 0 and 1) and $\alpha_0$ is the intracellular HBV DNA production rate; and an additional inhibitory effect of intracellular virus production rate g. κ and η are fixed to 0. g is fixed to 0.16 $d^{-1}$. B, D: We modelled the effect of treatment on degradation rate μ as follow: μ.κ and an additional inhibitory effect of intracellular virus production rate g where $\mu=0.01\ d^{-1}$ and ε and η are fixed to 0. g is fixed to 0.16 $d^{-1}$.

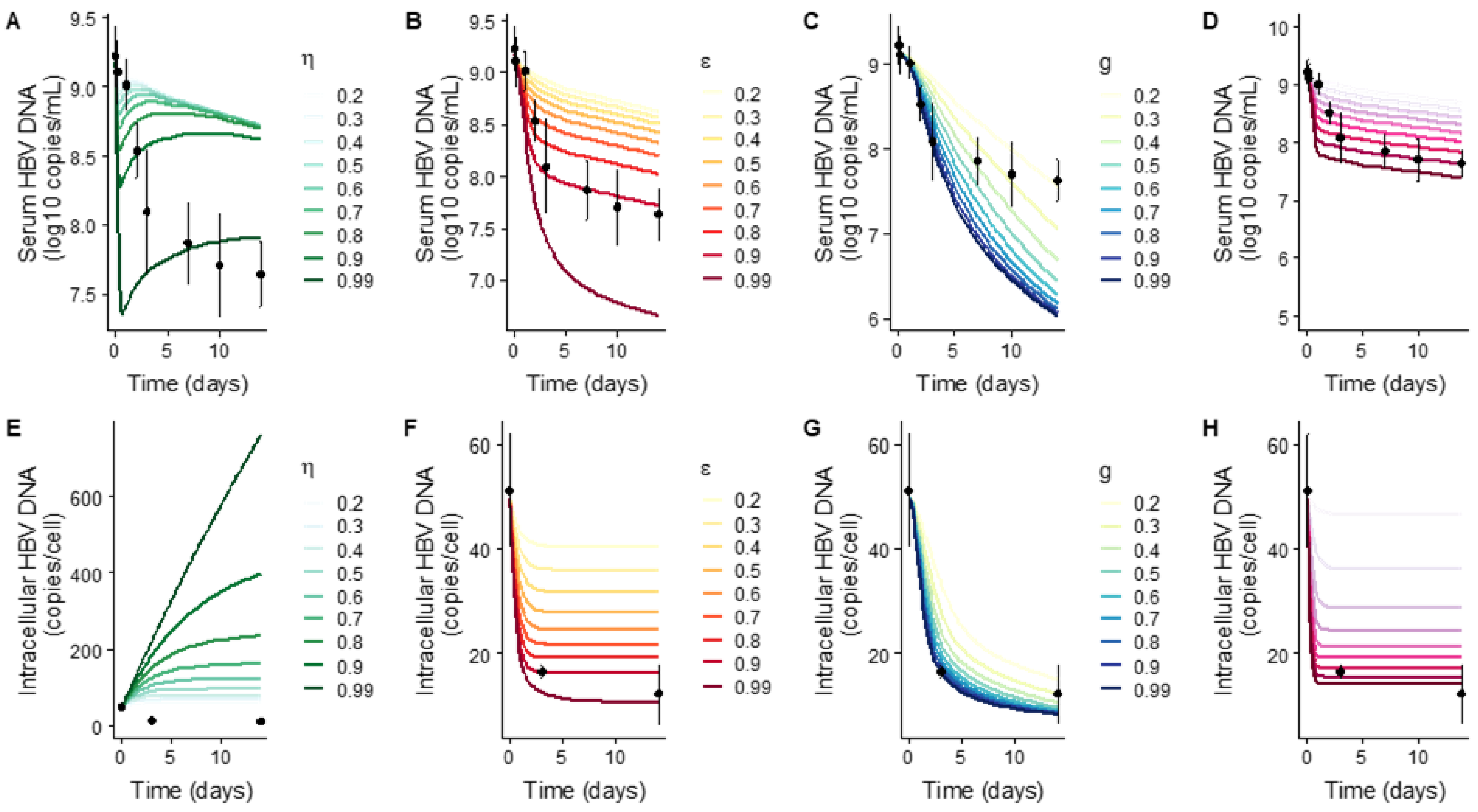


**Supplementary Figure 4: Simulation of serum (A, B, C, D) and intracellular (E, F, G, H) HBV DNA kinetics in pegIFN group and comparison with observed data using Eq. 1.**

**A, E:** We modelled the treatment effect as the inhibition of secretion as follow: $(1-\eta)\rho$ where $\eta$ is the drug effectiveness (varying between 0 and 1) and $\rho$ is the secretion rate. g, $\varepsilon$ and $\kappa$ are fixed to 0. **B, F:** We modelled the treatment effect as the intracellular HBV DNA production as follow: $(1-\varepsilon)\alpha_0$ where $\varepsilon$ is the drug effectiveness (varying between 0 and 1) and $\alpha_0$ is the intracellular HBV DNA production rate. g, $\kappa$ and $\eta$ are fixed to 0. **C, G:** We modelled the additional inhibitory effect on the intracellular HBV DNA production as follow: $\alpha_0 \exp(-g*t)$ where g is the additional inhibitory effect and $\alpha$ is the intracellular HBV DNA production rate. $\varepsilon$, $\eta$ and $\kappa$ are fixed to 0. **D, H:** We modelled the effect of treatment on degradation rate $\mu$ as follow: $\mu.\kappa$ where $\mu=0.01$ $d^{-1}$ and $\varepsilon$ and $\eta$ are fixed to 0. In all simulations model's parameters were fixed $I_0=3.10^8$ cells, $V_{,0}=$ 9.18 $log_{10}$ copies/mL, $D_0=50$ copies/cell, $\delta=0.01$ $d^{-1}$, $B_v=1.0$ mL, $c=11.8$ $d^{-1}$, $m=19.2$ copies/cell, $h=5.47$, $g=0$ $d^{-1}$ and $\mu=0.01$ $d^{-1}$. For $\delta(t)$, We used a step function to represent the biphasic decline. We used the average values of $s_1$ (0.094 $d^{-1}$), end of $s_1$ (5 d) and $s_2$ (0.054 $d^{-1}$) computed for the human albumin decline in mice from the IFN group.

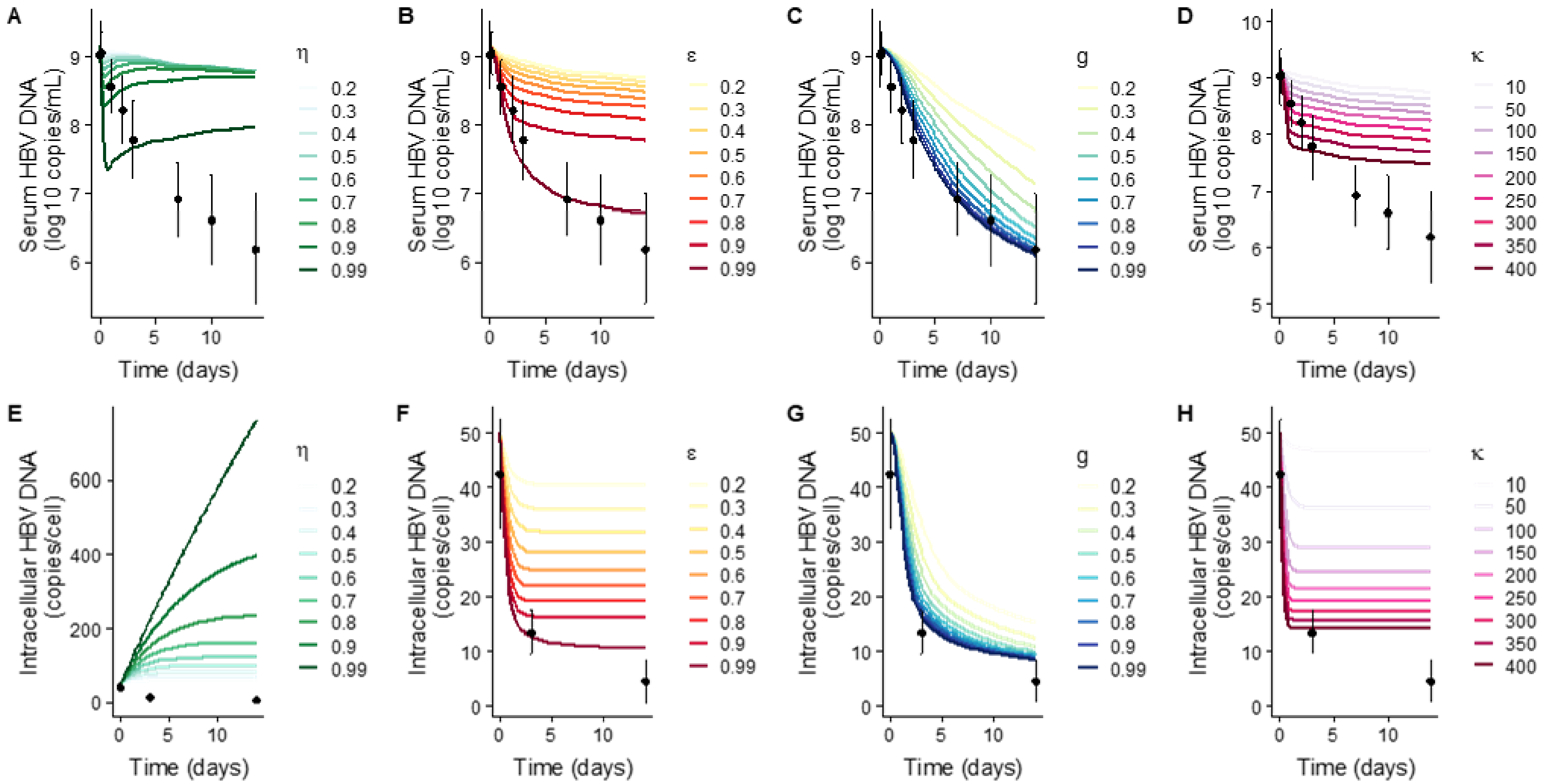


**Supplementary Figure 5: Simulation of serum (A, B, C, D) and intracellular (E, F, G, H) HBV DNA kinetics in LAM+IFN group and comparison with observed data using Eq. 1.**

**A, E:** We modelled the treatment effect as the inhibition of secretion as follow: $(1-\eta)\rho$ where $\eta$ is the drug effectiveness (varying between 0 and 1) and $\rho$ is the secretion rate. g, $\varepsilon$ and $\kappa$ are fixed to 0. **B, F**: We modelled the treatment effect as the intracellular HBV DNA production as follow: $(1-\varepsilon)\alpha$ where $\varepsilon$ is the drug effectiveness (varying between 0 and 1) and $\alpha$ is the intracellular HBV DNA production rate. g, $\kappa$ and $\eta$ are fixed to 0. **C, G:** We modelled the additional inhibitory effect on the intracellular HBV DNA production as follow: $\alpha_0 \exp(-g{*}t)$ where g is the additional inhibitory effect and $\alpha_0$ is the intracellular HBV DNA production rate. $\varepsilon$, $\eta$ and $\kappa$ are fixed to 0. **D, H**: We modelled the effect of treatment on degradation rate $\mu$ as follow: $\mu.\kappa$ where $\mu$=0.01 $d^{-1}$ and $\varepsilon$ and $\eta$ are fixed to 0. In all simulations model's parameters were fixed $I_0$=3.10$^{8}$ cells, $V_0$= 9.18 $\log_{10}$ copies/mL, $D_0$=50 copies/cell, $\delta$=0.01 $d^{-1}$, $B_v$ =1.0 mL, c=11.8 $d^{-1}$, m=19.2 copies/cell, h=5.47, g=0 $d^{-1}$ and $\mu$=0.01 $d^{-1}$. For $\delta(t)$, we used a step function to represent the biphasic decline. We used the average values of $s_1$ (0.088 $d^{-1}$), end of $s_1$ (6.44 d) and $s_2$ (0.031 $d^{-1}$) computed for the human albumin decline in mice from the PegIFN+LAM group.

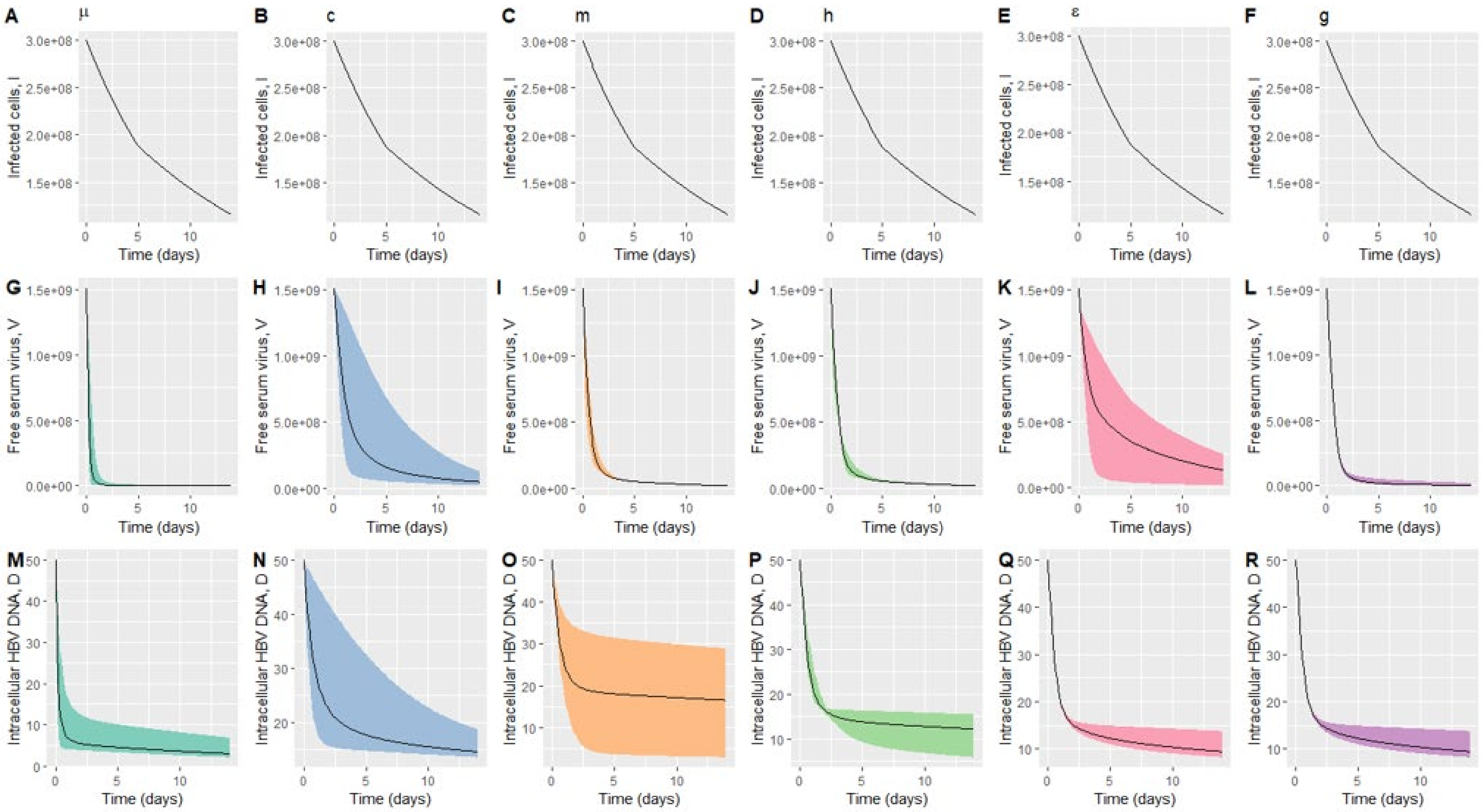


**Supplmentary Figure 6. Sensitivity of infected cells, I, (A-F), free serum virus, V, (H-M) and intracellualr HBV DNA, D, (O-T) to range of model parameters .**

Using the FME package in R3.3.2, we studied the practical identifiability of the model parameters at steady-state, as suggested by Brun et al. (3). We used the following ranges: for μ [0.001 – 10 $d^{-1}$] in panels A, G, M; for c [0.1-15 $d^{-1}$] in panels, B,H,N; for m [0-50 copies/cell] in panels C, I, O; for h [0.1 - 10] in panels D, J, P; for ε [0-1] in panels E, K, Q and for g [0-1 $d^{-1}$] in panels F, L, R. As expected, none of the parameters had an effect on infected cells (panels A-G). For all the other parameters, varying their values impacts free serum virus, V and/or intracellular HBV DNA, D kinetics. We found that collinearity was <20 only for models where μ was fixed.

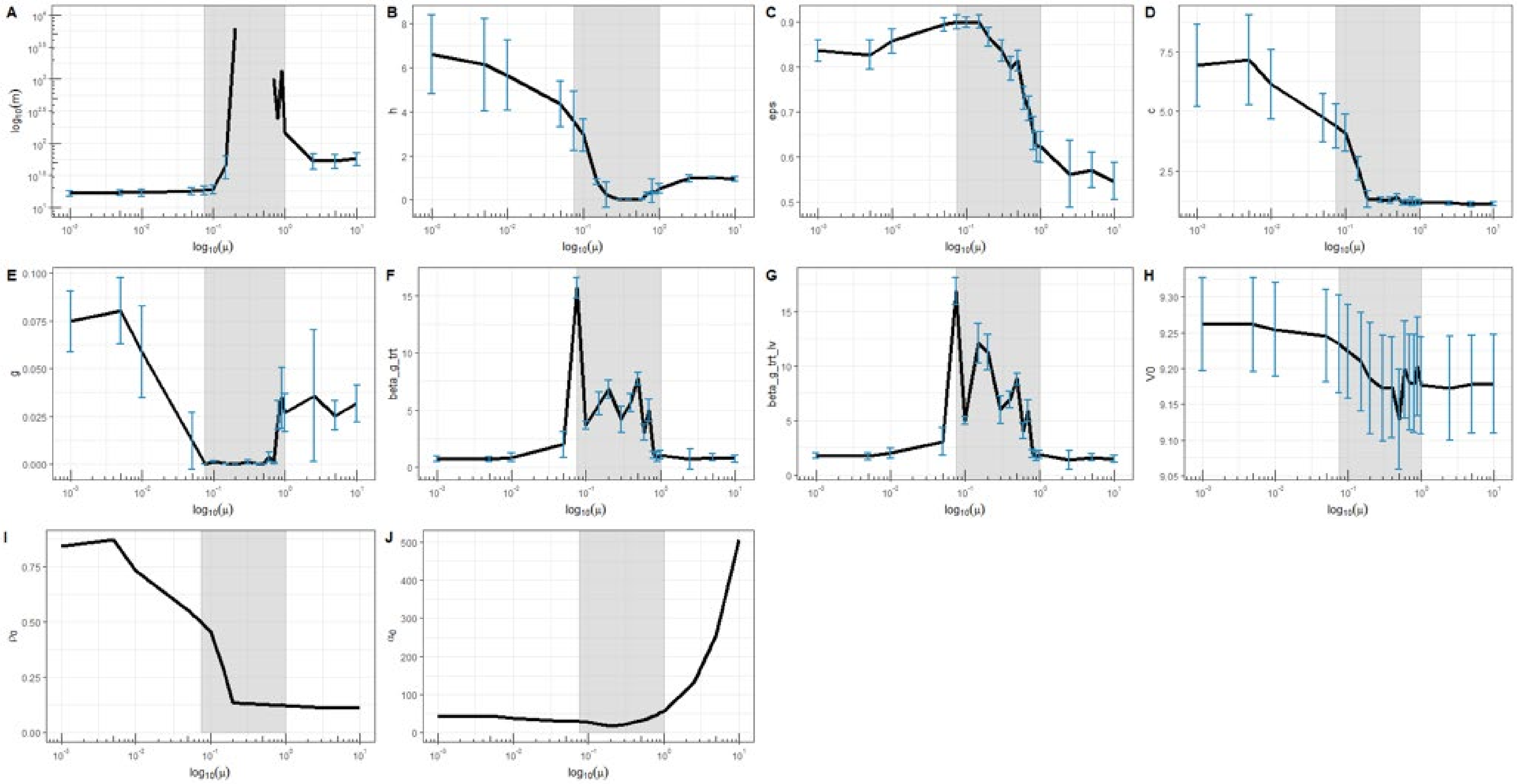

**Supplementary Figure 7. Parameter estimates for different values of pre-treatment intracellular virus clearance, μ, varying between 0.001 d$^{-1}$ and 10 d$^{-1}$.**

The grey interval where $\mu \in [0.075 - 1.00]$ d$^{-1}$ corresponds to the interval where the model did not converge and/or where parameters were poorly estimated. The blue error bars represent the interval: estimate ± standard error. Each box represents a parameter (A: m, B: h, C: ε, D: c, E: g, F: $\beta_{LAM}$, G: $\beta_{pegIFN+LAM}$, H: $V_0$, I: $\rho_0$ and J: $\alpha_0$). $\rho_0$ and $\alpha_0$ were computed from the other estimates.
